# Radiation induced toxicity in rectal epithelial stem cell contributes to acute radiation injury in rectum

**DOI:** 10.1101/2020.09.10.288696

**Authors:** Felipe Rodriguez Tirado, Payel Bhanja, Eduardo Castro-Nallar, Ximena Diaz Olea, Catalina Salamanca, Subhrajit Saha

**Affiliations:** Department of Radiation Oncology; Department of Cancer Biology, University of Kansas Medical Center, 3901 Rainbow Boulevard, Kansas City, Kansas 66160; Center for Bioinformatics and Integrative Biology, Facultad de Ciencias de la Vida, Universidad Andres Bello, Chile

**Author notes:** Correspondence should be addressed to: Subhrajit Saha PhD, Associate Professor, Department of Radiation Oncology, Department of Cancer Biology, The University of Kansas Medical Center, MS 4033, 3901 Rainbow Boulevard, Kansas City, Kansas 66160.

**Keywords:** Rectal stem cell, Pelvic irradiation, Crypt, mice

## Abstract

**Background:** Radiation induced rectal epithelial damage is a very common side effect of pelvic radiotherapy and often compromise the life quality and treatment outcome in patients with pelvic malignancies. Unlike small bowel and colon effect of radiation in rectal stem cells has not been explored extensively. Here we demonstrate that Lgr5 positive rectal stem cells are radiosensitive and organoid based transplantation of rectal stem cells mitigates radiation damage in rectum

**Methods:** C57Bl6 male mice (JAX) at 24 h was exposed to pelvic irradiation (PIR) to determine the radiation effect in pelvic epithelium. Effect PIR on Lgr5-positive rectal stem cells (RSCs) was determined in Lgr5-EGFP-Cre-ERT2 mice exposed to PIR. Effect PIR or clinically relevant fractionated PIR on regenerative response of Lgr5-positive RSCs was examined by lineage tracing assay using Lgr5-eGFP-IRES-CreERT2; Rosa26-CAG-tdTomato mice with tamoxifen administration to activate Cre recombinase and thereby marking the ISC and their respective progeny. Ex vivo three-dimensional organoid cultures were developed from Lgr5-EGFP-Cre-ERT2 mice. Organoid growth was determined by quantifying the budding crypt/total crypt ratio. Organoids from Lgr5-EGFP-ires-CreERT2-TdT mice were transplanted in C57Bl6 male mice exposed to PIR. Engraftment and repopulation of Lgr5-positive RSCs were determined after tamoxifen administration to activate Cre recombinase in recipient mice. Statistical analysis was performed using Log-rank (Mantel-Cox) test and paired two-tail *t* test.

**Result:** Exposure to pelvic irradiation significantly damaged rectal epithelium with the loss of Lgr5+ve rectal stem cells. Radio-sensitivity of rectal epithelium was also observed with exposure to clinically relevant fractionated pelvic irradiation. Regenerative capacity of Lgr5+ve rectal stem cells were compromised in response to fractionated pelvic irradiation. Ex-vivo organoid study demonstrated that Lgr5+ve rectal stem cells are sensitive to both single and fractionated radiation. Organoid based transplantation of Lgr5+ve rectal stem cells promote repair and regeneration of rectal epithelium.

**Conclusion:** Lgr5 positive rectal stem cells are radio-sensitive and contribute to radiation induced rectal epithelial toxicity. Transplantation of Lgr5 positive rectal stem cells mitigates radiation induced rectal injury and promote repair and regeneration process in rectum.

## Introduction

Rectal injury is a major limiting factor for definitive chemo-radiation therapy of pelvic malignancies, such as prostate cancer, bladder cancer or ovarian cancer, collectively 20-30% of all malignancies[1, 2]. Thus, effective doses of radiation and/or chemotherapy often cannot be administered, resulting in poor survival and early metastatic spread. Even with stereotactic radiosurgery targeting the tumor, the dose may be limited by an increase in acute rectal epithelial loss leading to acute bleeding followed by chronic proctitis[3]. Radiation proctitis is a common and debilitating consequence of radiation therapy induced damage to the rectal tissue characterized by acute mucosal loss, inflammation, followed by progressive tissue scarring leading to organ fibrosis. Currently there are no FDA-approved treatments that can be used to protect against radiation induced damage to the rectal epithelium. Previous research on radiation induced toxicity in small bowel demonstrated that radiation induced loss of intestinal stem cell is the major cause of mucosal damage[4-6]. Unlike radiation induced small bowel toxicity majority of the reports on radiation induced rectal injury described radiation induced proctitis /fibrosis[7, 8]. However, external beam pelvic radiotherapy very frequently causes acute injury such as rectal mucosal damage along with rectal hemorrhage/bleeding. To date not much been known about radiation induced toxicity in rectal epithelial stem cells.

Adult epithelial stem cells in small bowel are primarily located in crypt base and function as structural building block for repair and regeneration[4, 9]. Crypts present within the mouse small intestine have two types of stem cells. Bmi1 positive intestinal stem cells (ISCs) that are long-lived, label-retaining stem cells present at the +4 position of the crypt base. These Bmi1+ve ISCs interconvert with more rapidly proliferating LRG5+ve stem cells known as CBCs that express markers including Lgr5, Olfm4, Lrig1 and Ascl2[10-15]. Both in vivo mice model and ex-vivo organoid model it has been demonstrated that sensitivity of these stem cell are the determining factor for mucosal epithelial response to radiation exposure[4, 6]. In small intestine it has been reported that label-retaining stem cells present at the +4 position are more radio-resistant than CBCs and functions as a reserve stem cell pool.

In rectum the presence of similar stem cell system has not been reported. Moreover the involvement of these stem cells in regenerative response of rectal epithelium against radiation injury has not been studied well. An extensive literature shows that organoids derived from intestines recapitulate structural and functional characteristics of the tissue of origin. Organoids maintains basic crypt physiology [9] depending on stem cell survival and self-renewal. In the present study we have demonstrated that radiation induced loss of Lgr5+ve rectal stem cell is the major cause of rectal epithelial damage. In mice model as well as in ex vivo organoid model we have demonstrated that exposure to clinically relevant fractionated dose significantly reduce Lgr5+ve rectal stem cell along with epithelial regeneration. Rectal transplantation of rectal organoid in mice exposed to pelvic irradiation demonstrated engraftment of repopulating rectal stem cells and thereby repair and regeneration of rectal epithelium.

## Method

### Animals

Six to 10 week-old male C57BL6/J mice, Lgr5-eGFP-IRES-CreERT2 mice, B6.Cg-Gt(ROSA)26Sortm9(CAG-tdTomato)Hze/J mice and Gt(ROSA)26Sortm4(ACTB-tdTomato-EGFP)Luo/J mice were purchased from Jackson laboratories. For lineage tracing experiments, Lgr5-eGFP-IRES-CreERT2 mice were crossed with B6.Cg-Gt(ROSA)26Sortm9(CAG-tdTomato)Hze/J mice to generate mice Lgr5-eGFP-IRES-CreERT2; Rosa26-CAG-tdTomato heterozygote. The animals were maintained ad libitum and all studies were performed under the guidelines and protocols of the Institutional Animal Care and Use Committee of the University of Kansas Medical Center. All the animal experimental protocols were approved by the Institutional Animal Care and Use Committee of the University of Kansas Medical Center (ACUP number 2019-2487).

### Histology

The rectum of each animal was dissected, washed in PBS to remove intestinal contents and the rectum was fixed in 10% neutral-buffered formalin before paraffin embedding. Tissue was routinely processed and cut into 5?μm sections for haematoxylin and eosin and immunohistochemical staining. All haemotoxylin and eosin (HE) (Fisher Scientific, Pittsburgh, PA) staining was performed at the Pathology Core Facility in the KUMC Cancer Center.

To visualize rectal epithelial cell proliferation, each mouse was injected intraperitoneally with 120 mg kg^−1^ BrdU (Sigma-Aldrich, USA) 2 to 4 h before killing and rectum was collected for paraffin embedding and BrdU immunohistochemistry. Tissue sections were routinely deparaffinized and rehydrated through graded alcohols and incubated overnight at room temperature with a biotinylated monoclonal BrdU antibody (Zymed, South Francisco, CA). Nuclear staining was visualized using Streptavidin-peroxidase and diaminobenzidine (DAB) and samples were lightly counterstained with haematoxylin. Rectum from mice, not injected with BrdU, was used as a negative control. Digital photographs of rectal crypts were taken at high (× 40–60) magnification (Zeiss AxioHOME microscope) and crypt epithelial cells in intestinal sections were examined using ImageJ software and classified as BrdU positive if they grossly demonstrated brown-stained nuclei from DAB staining or as BrdU negative if they were blue stained nuclei.

### Determination of Crypt Depth

Crypt depth was independently and objectively analyzed and quantitated in a blind manner from coded digital photographs of crypts from HE-stained slides using ImageJ 1.37 software to measure the height in pixels from the bottom of the crypt to the top. This measurement in pixels was converted to length (in μm) by dividing with the following a conversion factor (1.46 pixels μm−1).

### Organoids culture

Rectum from C57BL/6 and Gt(ROSA)26Sortm4(ACTB-tdTomato-EGFP)Luo/J was used for Crypt isolation and development of *ex vivo* organoid culture. Tissue was washed gently in cold PBS (Calcium and Magnesium free) to clean fat and stool. Tissue was cut approximately into 3 mm pieces and washed with 10 ml of cold PBS using a serological pipette. The pieces were allowed to settle by gravity and supernatant was discarded. This washing step was repeated few times until clear supernatant was visible. Clean rectal tissue was then incubated in Gentle Cell Dissociation Reagent (Stemcell Technologies #07174) for 15 minutes at room temperature on a rocking platform (20 RPM). Once settled at the bottom of the tube, tissue pieces were re-suspended in sterile 0.1% BSA containing PBS solution and pipetted up and down three times to release villi portion (fraction one). This step was repeated four times to release rectal crypts from the tissue pieces. These four fractions were passed through 70 μm filters (BD Biosciences), and centrifuged at 275*g* for 5 min at 4 °C and single cells were discarded. Crypt pellet was resuspended in 10 ml cold (2-8^°^C) DMEM-F12 and centrifugated at 200 x g for five minutes. Isolated crypts were resuspended in cold Matrigel (Corning #356231) and cold media (StemCell Technologies IntestiCult^™^ #06005) in a 50:50 ratio. This was seeded as a dome in center of the wells of 48 well plate previously warmed a 37^°^C to solidify the Matrigel. After 15 min of incubation at 37^°^C when Matrigel was solid, culture media was added from the side of the well. Plates were maintained at 37^°^C with 5% CO_2_ with media change every 3 days.

### Irradiation procedure

Pelvic irradiation in single or multiple fractions was performed on anesthetized mice (intraperitoneal injection of Ketamine (87.5 mg/Kg) and Xylazine (12.5 mg/Kg) cocktail) using XenX (Xstrahl, Life Sciences)[6, 16]. A 3 cm area of the mice containing Pelvic region was irradiated (Fig.1A), thus shielding the upper and middle abdomen, thorax, head and neck as well as lower and upper extremities, protecting a significant portion of the bone marrow. The total irradiation time to deliver the intended dose was calculated with respect to dose rate, radiation field size and fractional depth dose to ensure accurate radiation dosimetry.

### Organoid Transplantation

Rectal organoids from Lgr5-eGFP-IRES-CreERT2; Rosa26-CAG-tdTomato mice were transplanted to C57Bl6 mice exposed to pelvic irradiation (24Gy single fraction). Around 150-200 organoids were transplanted in mice rectum through anus using a micropipette. Following transplantation recipient mice were treated with Tamoxifen to activate Cre recombinase in transplanted cells.

### *In vivo* Lineage tracing assay

Lgr5-eGFP-IRES-CreERT2 mice were crossed with B6.Cg-Gt(ROSA)26Sortm9(CAG-tdTomato)Hze/J mice (Jackson Laboratories)[17] to generate Lgr5-eGFP-IRES-CreERT2; Rosa26-CAG-tdTomato heterozygote. Involvement Lgr5 RSCs in rectal epithelial regeneration was examined by lineage tracing assay. Lineage tracing was induced by tamoxifen administration in Cre reporter mice to mark the RSCs and their respective tdT positive progeny. Adult mice were injected with tamoxifen (Sigma) (9 mg per 40 g of body weight, i.p.) to label Lgr5+ lineages.

### Immunocytochemistry

To validate the organoids derived from rectum expression of rectal epithelial specific markers such as Muc2, ChgA, CFTR were examined using immune-fluorescence analysis (supplement figure 1). In brief rectal organoids were permeabilized with 0.1% Triton X-100 (Sigma #9002-93-1) in PBS for 30 minutes at room temperature, washed twice with PBS and blocked with 5% normal goat serum (Life Technologies #50062Z) for 60 minutes. Organoids were incubated with primary antibodies (1:50 anti CFTR (Mouse IgM, Abcam #ab2784), 1:100 anti MUC2 (Rabbit IgG, Novus Biological #NBP1-31231) for overnight at 4^°^C followed by incubation with secondary antibodies (anti IgG Rabbit Alexa Fluor 488 (Invitrogen #A11008) 8 ug/ml, 1:50 anti IgM mouse Dylight 594 (Abcam #ab97009) for 5 hours at room temperature. For phospho-Histone H2A.X detection, organoids were stained with the Anti-phospho-Histone H2A.X (Ser139) Antibody (2 ug/ml) (Mouse IgG1, Millipore #16-202A, clone JBW301, FITC conjugate).

**Figure 1.**
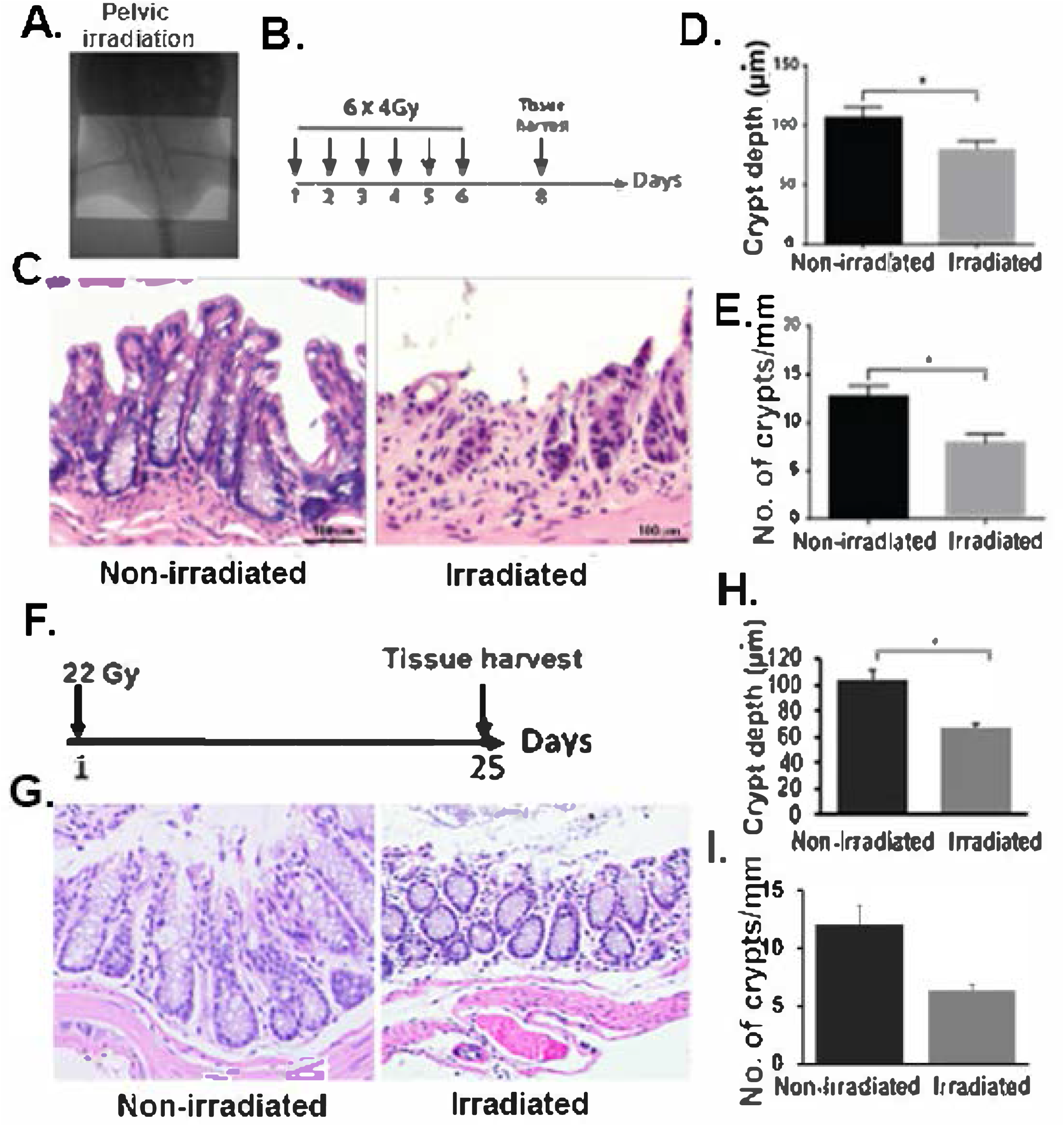
Radiation-induced damage to rectal tissue. **A**. Portal camera image demonstrating pelvic irradiation (PIR) exposure field. **B**. Schematic diagram demonstrating the timeline of fractionated pelvic irradiation. **C**. H&E stained representative cross-section of rectum. Non-irradiated mice showed a normal structure of crypts. However, mice exposed to fractionated pelvic radiation induced epithelial damage will loss of crypt. **D-E**. Histogram demonstrating crypt depth (C) and number of crypt per mm (D) on rectal tissue. Exposure to fractionated pelvic irradiation significantly reduces crypt depth (D) (n:3, p< 0.0061, Mann Whitney test) and number of crypt per mm (n:3 p-value: 0.0061, Mann Whitney test) on rectal tissue (**E)**. F. Schematic diagram demonstrating the timeline of single fraction pelvic irradiation. **G**. H&E stained representative cross-section of rectum. Non-irradiated mice showed a normal structure of crypts. However, mice exposed to pelvic radiation induced epithelial damage with loss of crypt. **H-I**. Histogram demonstrating mice exposed to pelvic irradiation decreases crypt depth (n:3, p<0.0048, Mann Whitney test) and crypt/mm (n:3 p< 0.0056, Mann Whitney test) on rectal tissue.

### ATP uptake-cell viability assay

This assay is designed for determining cell viability in 3D microtissue/organoid[18]. Cell Viability of the 3D organoid was measured using the CellTiter-Glo® 3D kit (Cat no.#. G968A, Promega, Madison-WI) using the manufacturer’s instruction. In this assay the reagent penetrates spheroids and using the lytic capacity accurately determines the viability by measuring the ATP uptake by the organoid. This 3D assay reagent measures ATP as an indicator of viability and generates a luminescent readout. In brief, Organoid were grown for 3-4 days in opaque-walled 96 well plates suitable for 3D cell culture. 48 hours after irradiation an equivalent volume of CellTiter-Glo_®_ 3D reagent was added in each well. The content of the plate was shaken for 5 minutes and luminescence was recorded 25 minutes after reagent addition. Different concentrations of ATP in water were plated at 100μl medium to generate an ATP standard curve. Luminescence of samples were compared to luminescence of standards to determine ATP detected by the CellTiter-Glo_®_ 3D Reagent in samples.

### RNAseq and Gene expression analyses

Flow-sorted Lgr5 positive and negative rectal epithelial cells were subjected to RNA sequencing. RNA samples were sent to BGI genomic center (China), paired-end cDNA libraries were prepared and sequenced using a BGISEQ-500RS. Raw reads were filtered and trimmed according to their quality score profiles as implemented in PrinSeq v0.20.4 (trimming at minimum quality of 20; low complexity reads (dust) = 32; no undetermined bases) [19]. High-quality reads were mapped against the mouse reference genome from NCBI (GCF_000001635.26_GRCm38.p6) using HISAT2 2.1.0 under default settings[20]. Mapped reads were converted to BAM format and sorted as in samtools[21]. We then assembled transcripts for each sample, estimated transcript abundances, and created table counts using StringTie 1.3.4d[22]. Transcript table counts were imported into the R environment using the tximport package. We performed a differential gene expression analysis using the negative binomial method with shrunk coefficients as implemented in the DESeq2 and package[23]. To explore enriched metabolic pathways, we performed a gene set analysis as implemented in the gage package[24], to then visualize such metabolic pathways in using the pathview package[25].

### Statistical Analysis

All images were analyzed using ImageJ software and graph were performed with GraphPad software. For histopathological and confocal images rectum regions were chosen at random for digital acquisition for quantitation. Number of animals used for all in vivo and ex vivo experiments were n=3 per group. A Mann Whitney test was used to determine significant differences between experimental conditions (P<0.05) with representative standard errors of the mean.

## Result

### Rectal epithelium is sensitive to clinically relevant fractionated radiation

Rectal epithelium structurally differs from small intestine and colon. Histopathological analysis demonstrated that rectum has elongated crypt with very minimal/short villus like structure as they do not have absorptive function. In the present study we have evaluated the effect of external beam fractionated radiation on mucosal epithelial structure of rectum. C57Bl6 mice were exposed to 6 fractions of 4Gy pelvic irradiation (Fig 1A-B) where each fraction was delivered in consecutive 6 days. At second day after last fraction of radiation rectum was subjected to histopathological analysis. H&E staining demonstrated significant loss of crypt like structure with reduction in crypt depth (Fig 1C-E). To develop an acute rectal injury model mice has been exposed to single fraction of pelvic irradiation (22Gy) (Fig 1F). Histopathological analysis demonstrated significant damage in epithelium with the loss of crypt and reduction in crypt depth (Fig 1G-I). These results clearly suggest that both fractionated radiotherapy nearly for a week as well as single fraction could promote acute injury in rectal epithelium loss of mucosal epithelial structure which may lead to severe pain/discomfort and blood loss.

### Radiation exposure reduces the regenerative capacity of Lgr5+ve RSCs

Mucosal epithelial loss is primarily consisting of stem cell loss along with dysregulated stromal signal[5]. In this study we have examined the effect of irradiation in rectal stem cell. To determine the effect of radiation in regenerative capacity of Lgr5+ve cells in rectum Lgr5-eGFP-IRES-CreERT2; Rosa26-CAG-tdTomato mice were exposed to high dose of single fraction of pelvic irradiation and then treated with single dose of Tamoxifen (Fig2A). In irradiated mice number of tdT positive cells which represents cells derived from Lgr5+ve RSCs significantly decreased compared to untreated control (Fig 2B-C).

**Figure 2.**
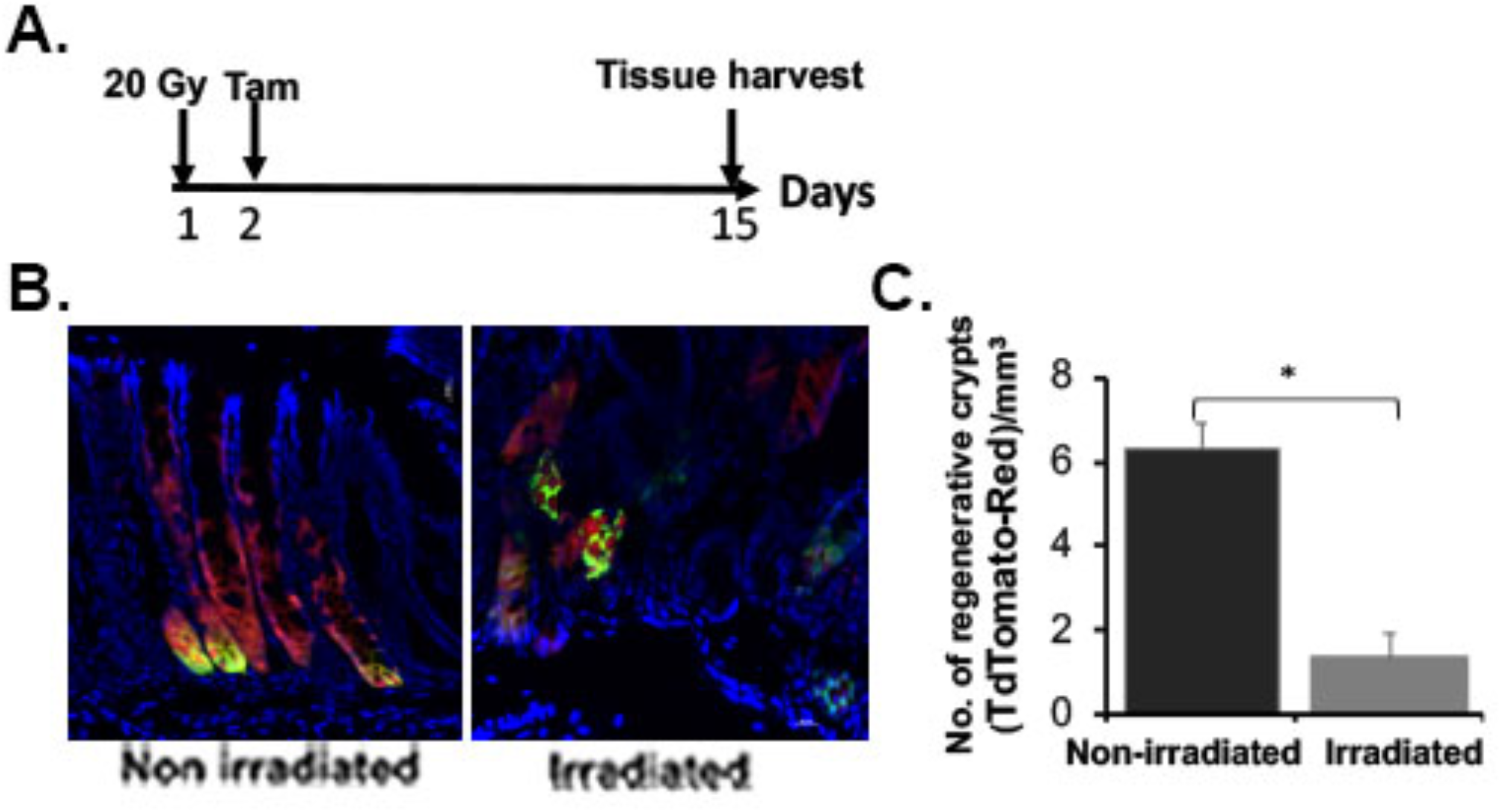
Pelvic irradiation reduces the regenerative response of LGR5+ stem cells in rectal crypts. **A**. Schematic representation of the treatment schema for lineage tracing assay in Lgr5-eGFP-IRES-CreERT2; Rosa26-CAG-tdTomato mice. **B**. Confocal microscopic images of the rectum section from Lgr5-eGFP-IRES-CreERT2; Rosa26-CAG-tdTomato mice. tdTomato (tdT)-positive cells are shown in red; Lgr5+ GFP+ cells are shown in green. Nuclei are stained with DAPI (blue). Marked expansion of tdT-positive red cells representing transit amplifying cells in crypt (regenerative crypt) were noted with un-irradiated rectal tissue compared to mice exposed to pelvic irradiation. **C**. The number of regenerative crypt significantly decreased in irradiated mice rectum compared to un-irradiated control (n=3; p<0.007).

Next we have examined the effect of fractionated radiation in regenerative capacity of rectal epithelial cells. Brdu analysis demonstrated presence of reduced number of Brdu+ve proliferating cells in rectal crypt from mice exposed to fractionated irradiation compared to unirradiated control (Fig 3A-C). To determine the regenerative capacity Lgr5+ve RSCs in response to fractionated irradiation Lgr5-eGFP-IRES-CreERT2; Rosa26-CAG-tdTomato mice were exposed to similar fractionated pelvic radiation dose regimen followed by Tamoxifen injection (Fig 3D). Confocal microscopic imaging demonstrated that significantly less number of tdTomato positive cells in irradiated group compared to un-irradiated control (Fig 3 E-F).

**Figure 3.**
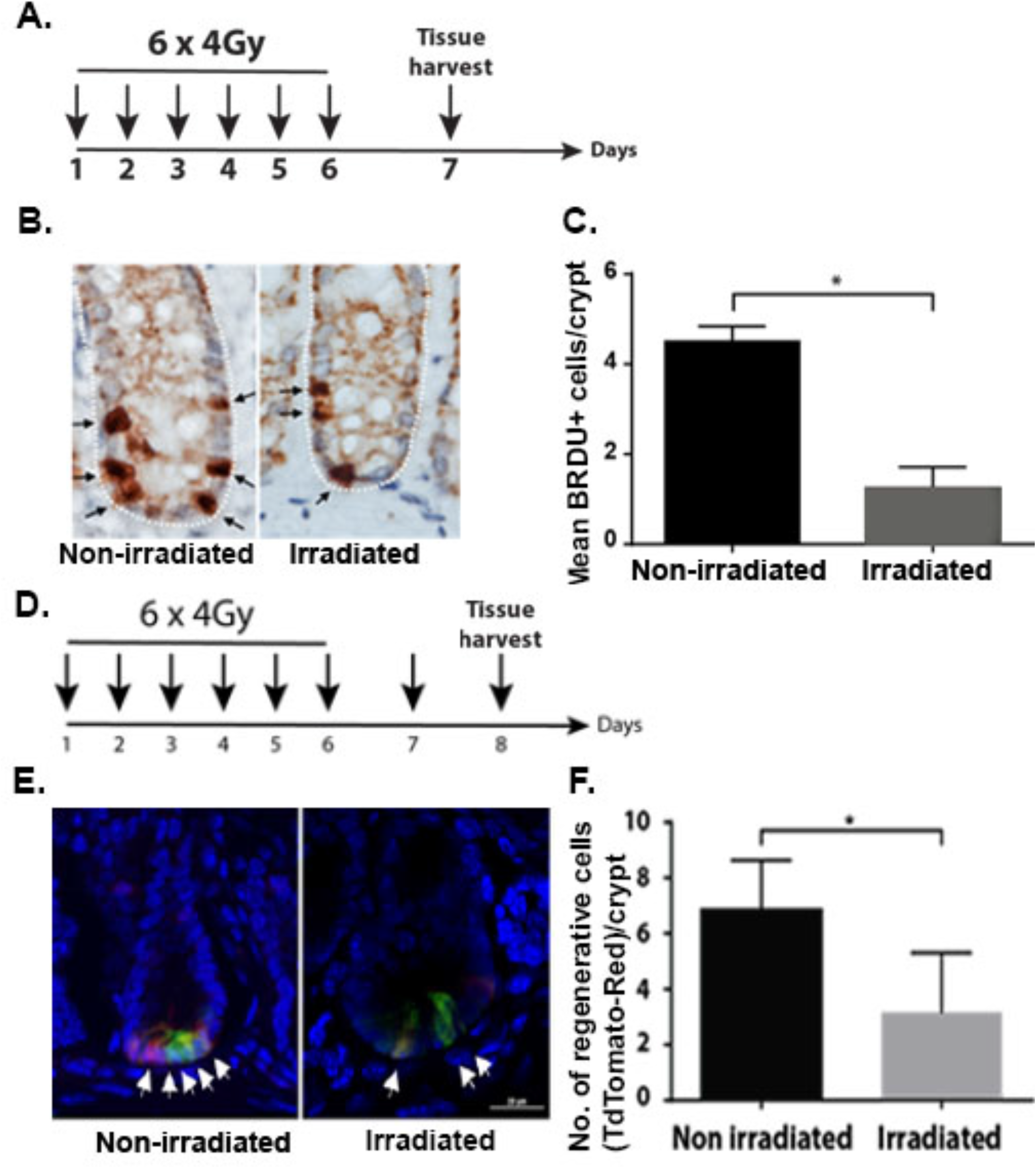
Fractionated pelvic irradiation impairs rectal epithelial regeneration. **A**. Schematic representation of the fractionated pelvic irradiation timeline. **B**. Representative Brdu immunohistochemistry of mice rectum section. Note the decrease in Brdu positive cells (stained brown, indicated with arrow). **C**. Histogram showing mean Brdu positive cells per crypt. Fractionated pelvic irradiation significantly reduce Brdu positive cells compared to un-irradiated control (P<0.0007, Mann Whitney test). **D**. Schematic representation of the fractionated pelvic irradiation and Tamoxifene treatment timeline. **E**. Confocal microscopic images of the rectum section from Lgr5-eGFP-IRES-CreERT2; Rosa26-CAG-tdTomato mice. tdTomato (tdT)-positive cells are shown in red; Lgr5+ GFP+ cells are shown in green. Nuclei are stained with DAPI (blue). Marked expansion of tdT-positive red cells representing transit amplifying cells (regenerative cells) in crypt were noted with un-irradiated rectal tissue compared to mice exposed to pelvic irradiation. **F**. The number of regenerative cells reduced significantly in irradiated mice rectum compared to un-irradiated mice (p<0.0381, Mann Whitney test).

### Fractionated irradiation inhibits rectal organoid growth

Intestinal organoids primarily from small intestinal and colonic crypts has been used extensively to study radiation effect[4, 6, 9, 16, 26, 27]. Organoids has also been considered as a model system to determine radiosensitivity for intestinal stem cells[27-29]. In this study for the first time we have used organoids from mice rectum to study radiation toxicity. Organoids in response to irradiation demonstrated significant decrease in survival with radiation dose dependent manner (Fig 4 A-B). We have also determined the effect of irradiation on overall survival of these organoids by using ATP assay. Irradiation reduces viability of these organoids in a dose dependent manner (Fig 4 C). To determine the effect of clinically relevant fractionated doses of radiation organoids were exposed to graded doses (2-8 Gy) of fractionated irradiation according to schedule described in Fig 5A. Representative bright-field images demonstrated lethality in organoids when exposed to fractionated irradiation (2Gy x 4) (Fig 5B). Organoids demonstrated complete loss in budding structure and eventually manifested as cloud of cell debris at day 9-11 post irradiation considering the days from first fraction of radiation (Fig 5B). Analysis of Feret diameter (Fig 5C), total organoid count (Fig 5D), percent budding organoids (Fig 5E) and viability percentage (Fig 5F) clearly demonstrated a dose response with graded doses of fractionated irradiation. Feret diameter is used to determine the organoid size[30]. Organoids exposed to irradiation clearly shows reduction in organoid size. Total organoid count also reduced significantly with irradiation suggesting increase death of organoids with a dose dependent manner. Quantification of budding organoids showed significant decrease in percentage of budding organoid in respect of total organoid count suggesting impaired growth and proliferation. These observations therefore suggest that rectal organoids which primarily depends on rectal stem cell for their survival are sensitive to fractionated irradiation. Moreover consistent dose dependent response of rectal organoids against radiation suggests that ex-vivo organoid platform could be used for radiation bio-dosimetry.

**Figure 4.**
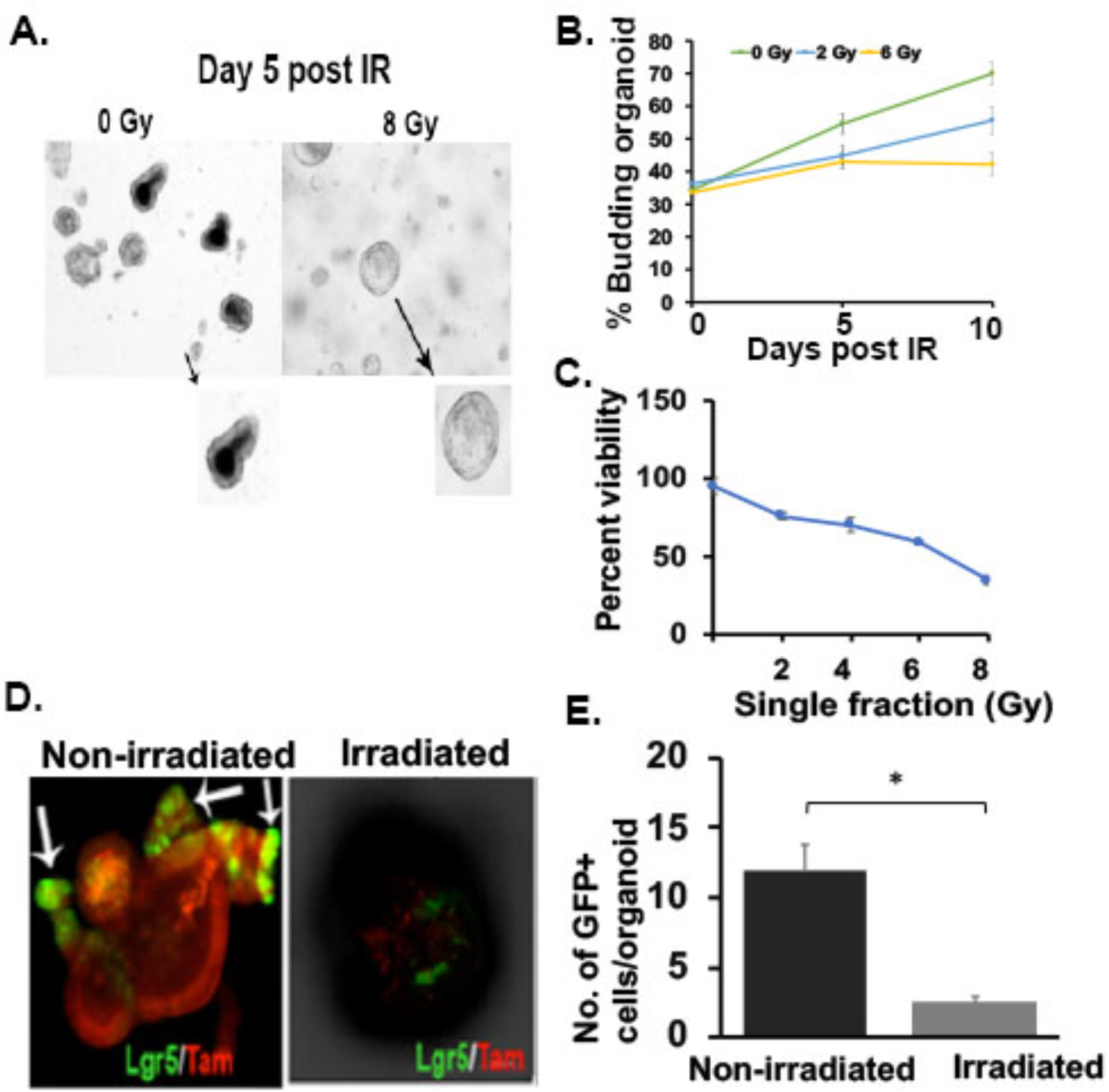
Effect of irradiation on rectal organoids. **A-B**. Microscopic image (phase contrast) of rectal organoids along with histogram of budding crypt/total crypt ratio demonstrating that irradiation impaired the organoid growth compared to un-irradiated control (2Gy *p<0.004, 6Gy *p<0.006). Microscopic image with 10X (indicated with arrow) and 40X magnification demonstrated loss of budding crypt in irradiated organoids. **C**. Survival assay (ATP uptake assay) demonstrated significant reduction of organoid survival in a dose dependent manner. **D**. Confocal microscopic images of organoids developed from Lgr5-EGFP-CRE-ERT2; R26-ACTB-tdTomato-EGFP mice demonstrated loss of Lgr5+ve cells (Green) in budding crypt from irradiated organoids compared to un-irradiated control. tdTomato is constitutively expressed in these mice as membrane bound protein therefore allows better visualization of cellular morphology. **E**. Histogram demonstrating significant decrease in Lgr5 +ve cells in irradiated rectal organoids compared to un-irradiated control (p<0.0004).

**Figure 5.**
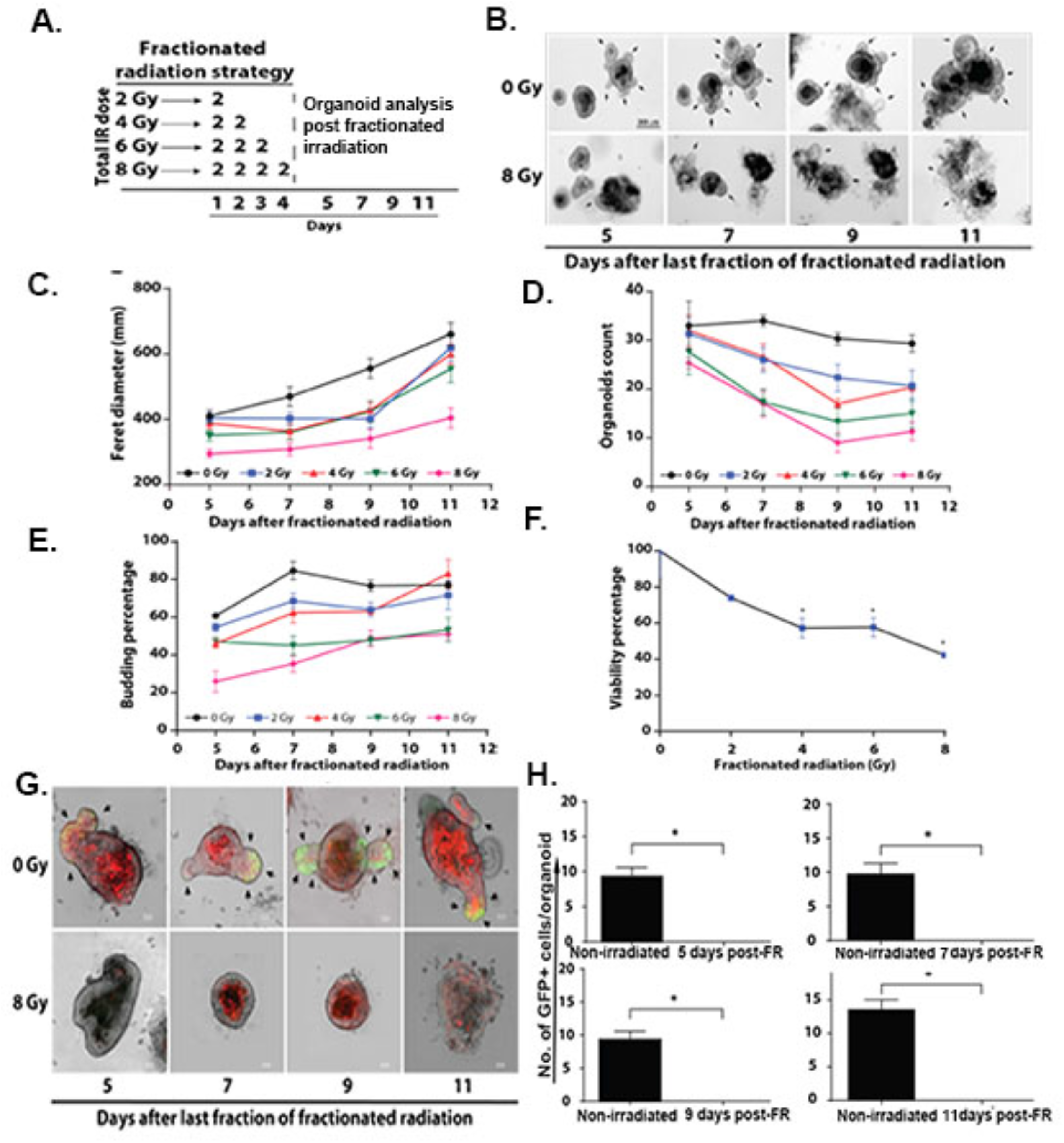
Effect of fractionated radiation on rectal organoids. **A**. Schematic representation of the fractionated irradiation to organoid. **B**. Microscopic image (phase contrast) of rectal organoids. **C-E**. Significant decrease in physical parameters such as (C) Feret diameter (8Gy vs 0Gy (p< 0.0001) at day 8 post IR) (D) organoid count (8Gy vs 0Gy (p< 0.0028) at day 8 post IR) and (E) budding percentage (8Gy vs 0Gy (0.0094) at day 8 post IR) was observed in a dose dependent manner. **F**. Significant decrease in organoid viability was observed in a dose dependent manner (8Gy vs 0Gy (0.0028) Mann Whitney test). **G**. Confocal microscopic images of organoids developed from Lgr5-EGFP-CRE-ERT2; R26-ACTB-tdTomato-EGFP mice demonstrated loss of Lgr5+ve cells (Green) in budding crypt from irradiated organoids compared to un-irradiated control. tdTomato is constitutively expressed in these mice as membrane bound protein therefore allows better visualization of cellular morphology. **H**. Histogram demonstrating significant decrease in Lgr5 +ve cells in irradiated rectal organoids compared to un-irradiated control at different time point post irradiation (p< 0.0079, day 11 post IR).

### Rectal stem cells in organoid is sensitive to irradiation

To determine the radiosensitivity of RSCs we have developed organoids from Lgr5-eGFP-IRES-CreERT2 mice and exposed to irradiation. Confocal microscopic imaging of un-irradiated organoids demonstrated the presence of Lgr5+ve cells at the tip of the budding crypt like structure. However, in response to single (Fig 4C-D) or multiple fraction of irradiation (Fig 5G-H) (2Gy ⨯ 4) demonstrated loss of Lgr5+ve cells with significant damage in overall organoid structure. Within days after last fraction of irradiation organoids demonstrated complete lethality as organoid structure end up as cloud of cell debris. These time course studies clearly suggest that radiation induced loss of Lgr5+ve RSCs is the earlier event which results complete inhibition of organoid growth and proliferation and eventual lethality.

### Radiation induces DNA damage in rectal organoids

In response to DNA double-strand break (DSB) H2AX the histone H2A variant becomes phosphorylated at serine 139. The phosphorylated form of H2AX is also known as γH2AX. To determine DNA damage response in irradiated rectal organoids number of γH2AX foci was quantified. Organoids stained with anti-γH2AX antibody demonstrated a significant increase in number of γH2AX foci following exposure to fractionated irradiation in a dose dependent manner (Fig 6A-B).

**Figure 6.**
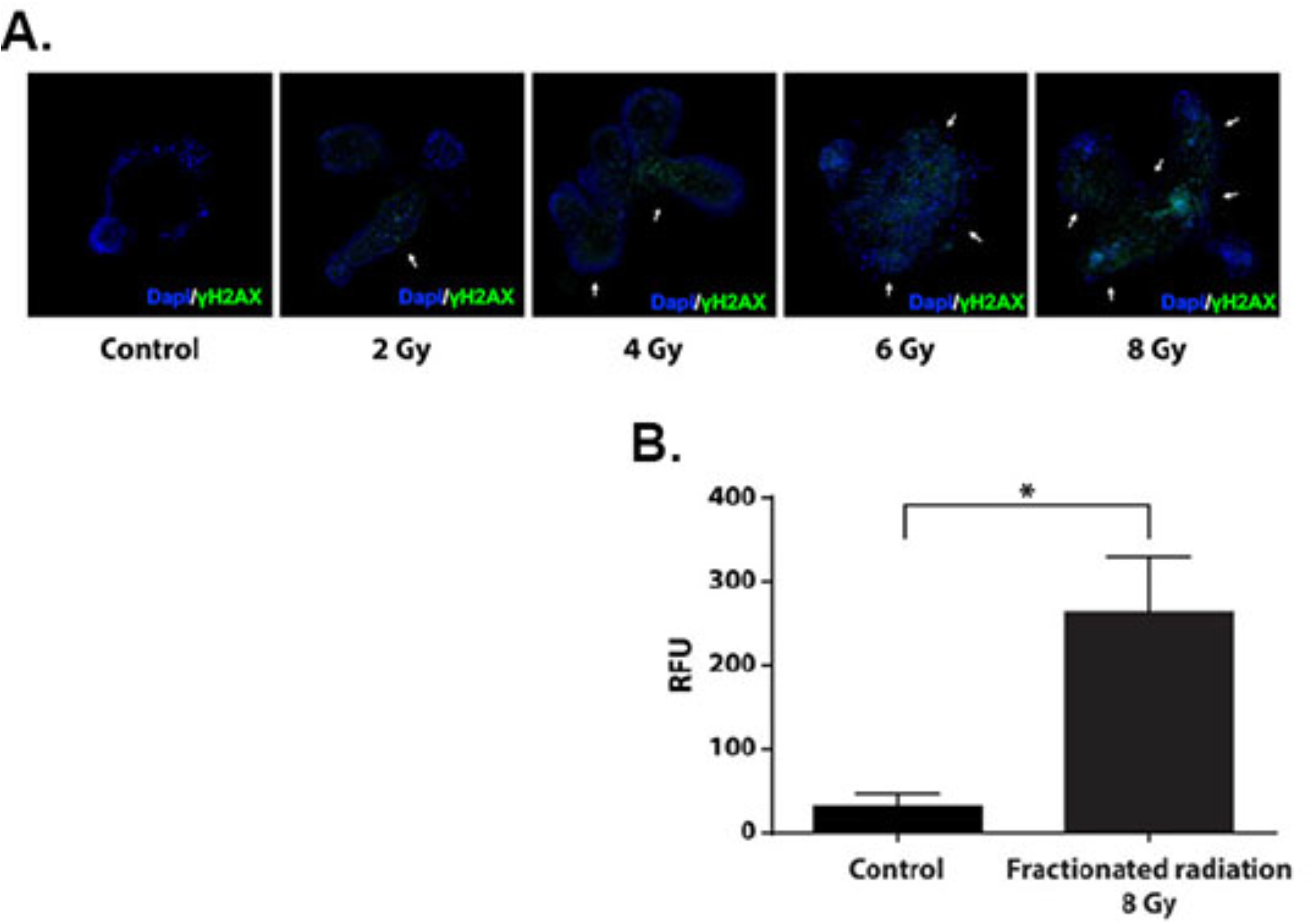
Radiation induce phosphorylation of histone H2A.X. **A**. Confocal microscopic images of rectal organoids exposed to 0, 2, 4, 6 and 8 Gy of fractionated radiation. Increase in Histone phosphorylation (γH2A.X green color) was observed in a dose dependent manner. **B**. Histogram demonstrating significant increase in Relative Fluorescence Units (RFU) representing immune-fluorescence signal in organoids exposed to 8 Gy of fractionated radiation, (n=4, p-value<0.05. Mann Whitney test) compared to un-irradiated control.

### Transplantation of rectal organoids mitigate radiation injury in rectum

Both single or multiple fraction radiation demonstrated significant loss of RSCs with impaired regeneration of rectal epithelium. Therefore, restitution of Lgr5 RSCs is critical for repair and regeneration. Transplanting organoid cells as a regenerative therapy has been used against degenerative diseases in intestine[31-33]. However, organoid based transplantation to mitigate radiation injury has not been examined in pre-clinical and clinical level. In the current manuscript ex-vivo rectal organoids grown from Lgr5-eGFP-IRES-CreERT2; Rosa26-CAG-tdTomato mice were transplanted to C57Bl6 mice exposed to pelvic irradiation (24Gy single fraction) (Fig 7A). Presence of Lgr5+GFP+tdT+ were observed in C57Bl6 mice rectum receiving organoid transplant (Fig 7B-C). Transplantation of organoids from Lgr5-eGFP-IRES-CreERT2; Rosa26-CAG-tdTomato having inducible Cre recombinase demonstrated presence of Lgr5+ cells derived tdT positive daughter cells in recipient C57Bl6 mice after Tamoxifene treatment (Fig 7B-C) suggesting regenerative response of RSCs. Histological analysis demonstrated loss of crypt and normal epithelial structure in irradiated rectum (Fig 7D). However, mice receiving organoid transplant post irradiation demonstrated presence of crypt with normal epithelial structure (Fig 7D). These results clearly suggest that organoid transplantation can induce repair and regeneration of irradiated rectal epithelium with the engraftment of regenerative RSCs.

**Figure 7.**
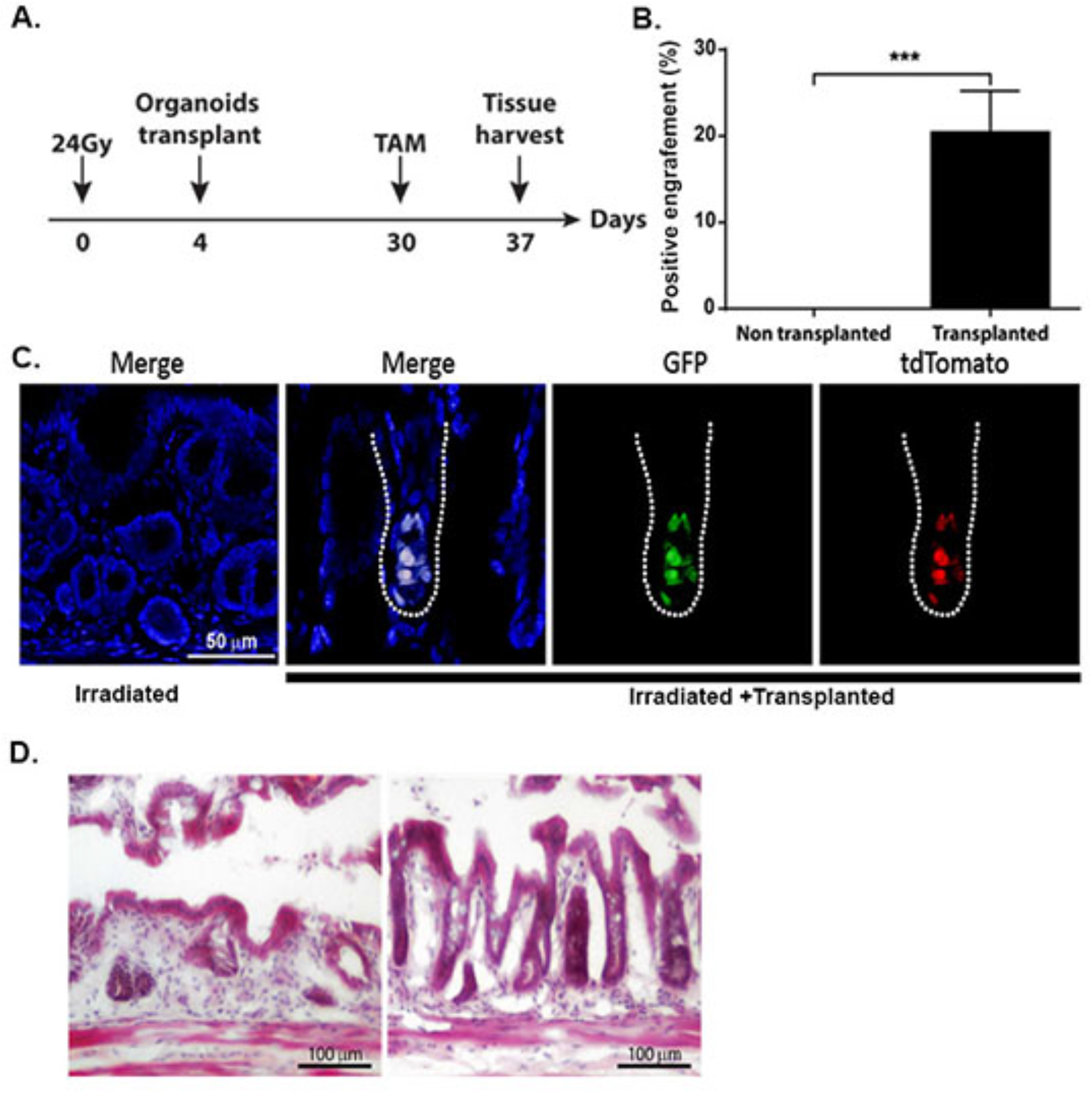
Transplantation of rectal organoids. **A**. Schematic diagram demonstrating pelvic irradiation and organoid transplantation time line. **B**. Histogram demonstrating significant presence of tdT cell containing microscopic fields (positive fields) in transplanted mice rectal epithelium compared to non-transplanted control (p<0.0005, Mann Whitney test n=4 mice (5-7 fields were counted per mice). **C**. Confocal microscopic image of rectal epithelium receiving transplanted organoids. Please note presence of GFP+ve (green) and tdTomato (red) cells in transplanted rectum suggesting presence of transplanted cells from Lgr5-eGFP-IRES-CreERT2; Rosa26-CAG-tdTomato mice rectal organoid. Merged figure demonstrated co-expression of Lgr5+GFP and reporter tdTomato. **D**. HE staining demonstrated rectal epithelial repair in transplanted mice compared to non-transplanted control.

### LGR5 positive cells expresses distinct stem cell marker in rectal crypts

Although ISCs were well characterized in small intestine and colon their rectal counterpart has not been studied well. LGR5 is a classic stem cell marker for intestinal stem cells localized in the bottom of crypts in small bowel and colon. These cells also express several other stem cell markers such as prominin1, musashi, Ascl2, OLMF4[34]. In the current report, using Lgr5-Cre-ERT-GFP mice we have already shown that Lgr5+ve cells were present in crypt bottom of rectum and responsible for regenerative response of rectal epithelium. To determine the presence of other known stem cell marker we have performed RNA seq analysis of sorted Lgr5+ve cells and Lgr5-ve cells (Fig 8A). We were able to identify 801 genes differentially expressed between these two populations (Fig 8B). Among these genes Lgr5 positive cells also expresses a series of known stem cell markers previously identified in intestines and other tissues. However, we have identified 9 markers which exclusively expressed in rectal Lgr5 positive cells compared to same cell types in small and large intestine (Fig 8C). Pathway based analysis suggested presence of cell cycle, DNA repair and recombination related genes in Lgr5+ve cells (Fig 8D).

**Figure 8.**
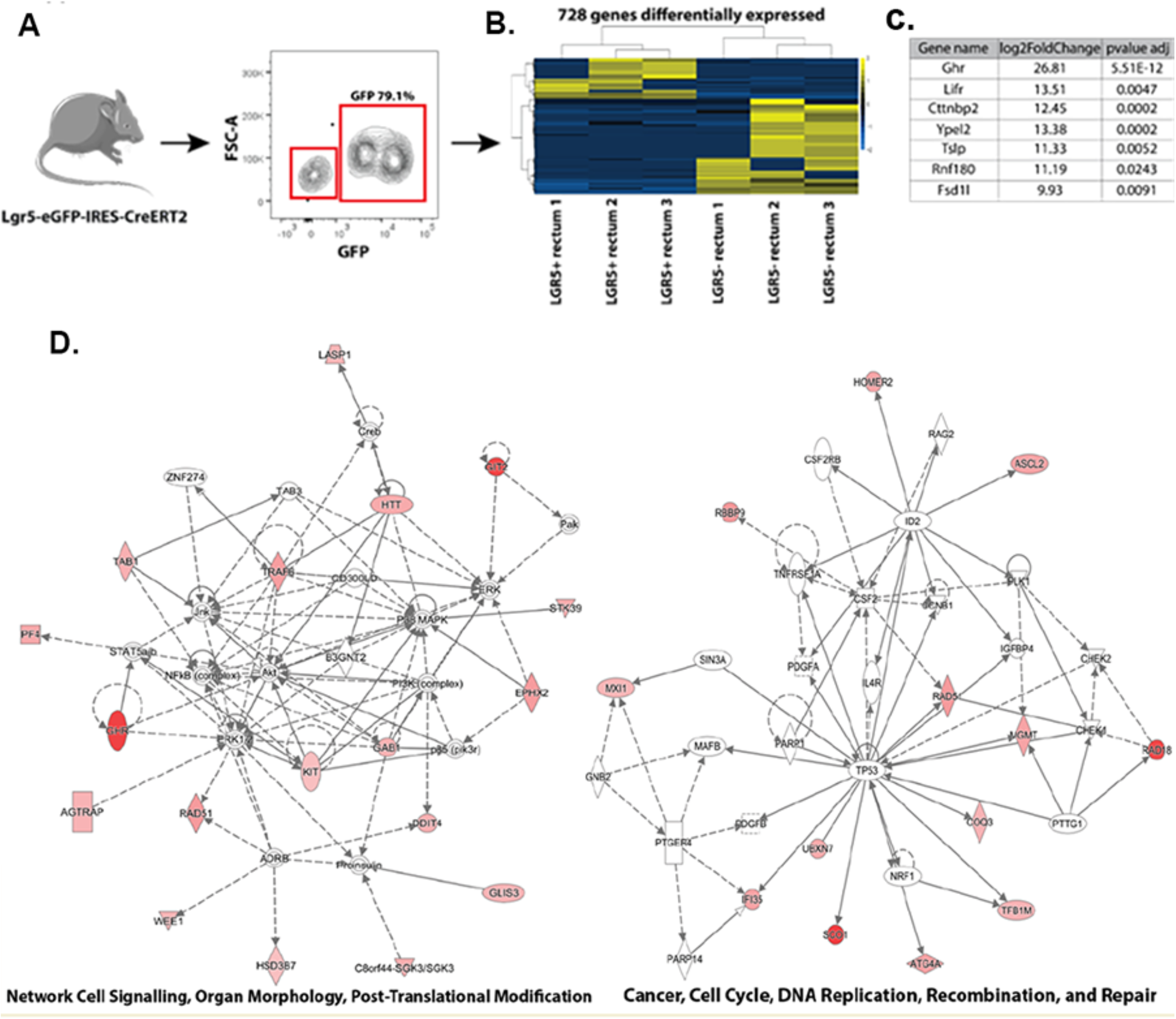
RNA seq analysis of Lgr5+ve and negative cells. LGR5+ and LGR5-cells derived from rectal crypts were sorted and sequenced to determine differential gene expression. **A**. Contour plot of flow-sorted LGR5+ (GFP+ve) and LGR5-ve rectal epithelial cells. **B**. The heat map showed 728 genes differentially expressed genes between LGR5+ and LGR5-cells derived from rectal tissue. **C**. List of stem cell marker exclusively expressed in rectum. (p-value adjusted cutoff of 0.05). These markers do not express in any other part of intestine. **D**. Ingenuity Pathway Analysis show genes up-regulated were principally associated with Cell signaling, post-translation modification, and organ morphology network and to Cell Cycle, DNA replication, Recombination and repair.

## Discussion

Results of the current study both in mice model and ex-vivo organoid model indicate that Lgr5+ve rectal stem cell is sensitive to acute radiation injury. We have generated an in vitro radiation sensitivity assay in organoid model. Findings of this study provide strong evidence in support of our in vivo data that clinically relevant fractionated irradiation cause acute loss of rectal stem cell in rectal epithelium which leads to rectal epithelial damage. Organoid based studies clearly demonstrated a dose dependent effect which is consistent both in multi and single fraction of irradiation. Current literatures primarily focused on the radiation induced rectal fibrosis which is late effect of radiation toxicity[7]. However, it is very critical to analyze the initial cause of onset of fibrosis. Our study demonstrates that clinical doses of pelvic irradiation impaired the rectal epithelial homeostasis and regeneration. Previous reports suggested loss of mucosal epithelium may promote repopulation of inflammatory cells and later fibrotic tissue[35]. Unlike small intestine and colon characterization of rectal stem cell has not been studied extensively. Our previous reports have demonstrated that Lgr5+ve stem cells in small intestine is radiosensitive and disappear within 72 hours after lethal dose of irradiation[4-6]. We have also shown that therapeutic intervention will only be successful to mitigate radiation induced acute mucosal damage if applied within 72 hrs post irradiation when endogenous Lgr5+ve stem cells are present[5]. Failure of mitigation in later time points clearly suggest that presence of Lgr5+ve stem cells are critical for rebuilding process. In another study from our group demonstrated that small molecule mediated activation Lgr5+ve stem cells in small intestine promote repair and regeneration[6]. In the present study we have shown that Lgr5+ve cells functions as stem cells in rectum and participate epithelial homeostasis and regeneration. Radiation induced depletion of these stem cells inhibits the regenerative process. Lineage tracing assay in mice demonstrated that in response to fractionated irradiation regenerative potential of these Lgr5+ve cells compromised significantly.

Radiation toxicity primarily induced in the cell by DNA damage. Our study demonstrates that fractionated doses of irradiation induces the DNA damage in organoid in dose dependent manner. Delivery of irradiation in multiple fraction with interval keeps level of DNA damage which promotes lethality in stem cells resulting organoid death. These results provide a mechanistic evidence of rectal epithelial damage when exposed to clinically relevant fractionated pelvic radiotherapy. To mitigate radiation induced rectal epithelial damage transplantation of rectal organoids enriched in rectal stem cells demonstrated repair and regenerative effect. Transplantation of organoids from Lgr5-eGFP-IRES-CreERT2; Rosa26-CAG-tdTomato having inducible Cre recombinase demonstrated presence of tdT+ cells in host rectal epithelium suggesting proliferation of Lgr5+ve cells. Histopathological evidences of rectal epithelial repair also suggested that organoid transplantation not only induce repair process by engraftment/cell replacement but may also release paracrine signals to induce the regenerative response of host stem/progenitor cells in irradiated rectum.

Previous reports showed that Lgr5+ cells in small intestine and colon shares other stem cell markers[34]. Our RNA seq data demonstrated that Lgr5+ cells in rectum shares a different set of stem cell markers compared to small intestine and colon. This data suggests that Lgr5+ cells in rectum may be different compared to other part of intestine and needs further investigation for their molecular and functional characterization. Previous reports demonstrated the presence of reserve and active pool of stem cells in small and large intestine which also differs in radio-sensitivity. It is possible that there could be a reserve stem cell pool in rectum such as Krt-19+ve cells[36] observed in small intestine or colon. Further studies are needed to determine the presence of other stem cell pool.

## Conclusion

In conclusion, the present study has demonstrated that Lgr5+ve RSCs are radiosensitive and it is critical to rescue these cells for repair and regeneration of rectal epithelium. Therefore, to ameliorate the acute toxicity in rectum in cancer patients undergoing pelvic radiotherapy a RSC targeted therapy is needed.

## List of abbreviations

ISC: Intestinal stem cells
RSC: Rectal stem cells
PBI: Partial body irradiation
HE: Hematoxylene and eosin

## Consent for publication

Not applicable.

## Availability of data and material

The authors declare that all data supporting the findings of this study are available within the article and its Supplementary Information files or from the corresponding author on reasonable request.

## Author Contribution

F.T. Conceived and designed the experiments, performed the experiments, analyzed the data, wrote the paper; P.B. Conceived and designed the experiments, performed the experiments; E.C.N: analyzed the data, X.D.O: performed the experiments; C.S: performed the experiments; S.S Contributed reagents/materials/analysis tools, conceived and designed the experiments, wrote the paper.

All authors read and approved the final manuscript.

## Acknowledgements

The authors acknowledge the histopathology core facility, confocal microscopy core facility of KUMC for their help and support.

