## SUPPLEMENT FIG 1 for "Radiation induced toxicity in rectal epithelial stem cell contributes to acute radiation injury in rectum"

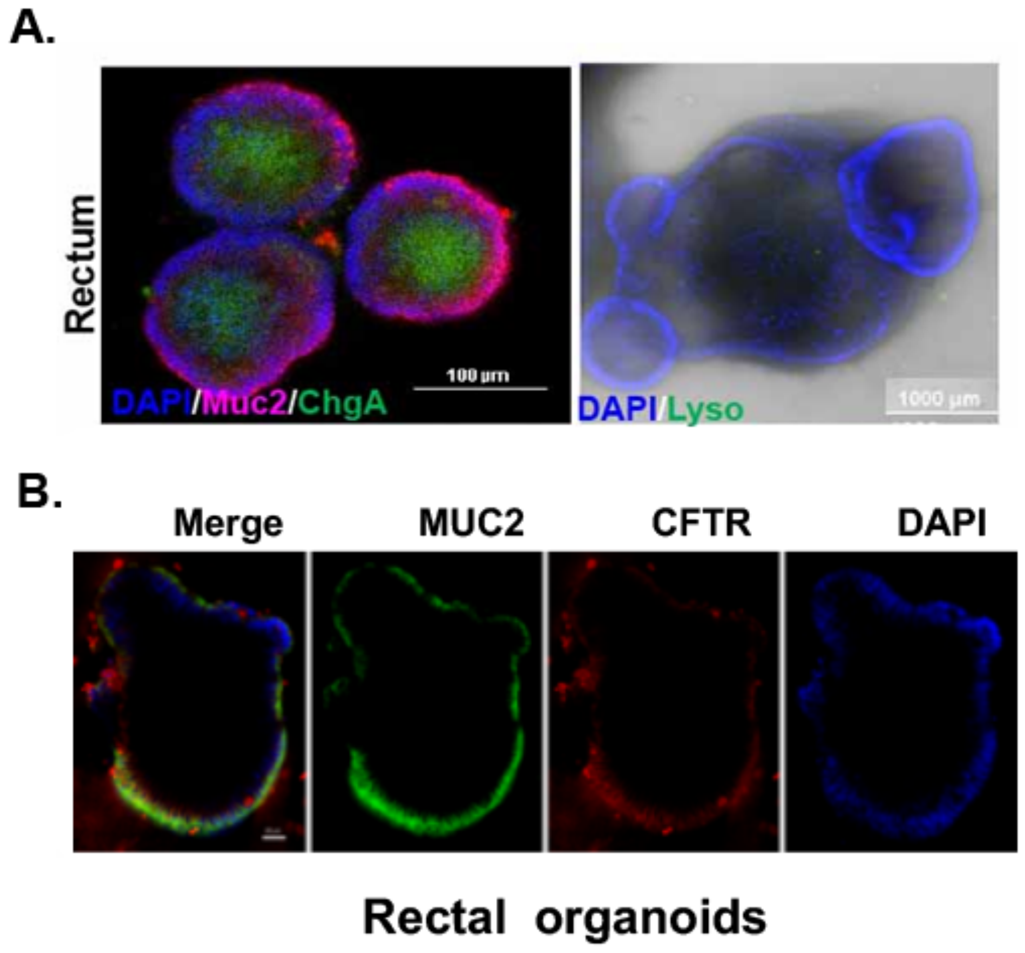


Supplement Figure 1

Fig1. Confocal microscopic image of the rectal organoid expressing markers predominant in rectal epithelium.

A) Representative image of mice rectal organoid co-expressing Muc2 (goblet cell marker) and ChgA (ChromograninA-enteroendocrine cell marker) (Left panel). Please note absence of paneth cell marker -Lysozyme expression predominantly expressed in small bowel and colon (right panel). B) Representative image of mice rectal organoid co-expressing Muc2 (goblet cell marker) and CFTR (cystic fibrosis transmembrane conductance regulator) protein specifically predominant in rectal epithelium.
